# Distinct RNA-RNA spatial interaction sub-networks of transcription factors in hepatic cells

**DOI:** 10.64898/2026.09.13.751294

**Authors:** Yuan Yang, Jun Xu, Hengdong He, Yuanling Tang, Xingyu Li, Yunhan Jing, Chen Wang, Yan Ding, Mingzhou Li, Qianzi Tang

**Affiliations:** State Key Laboratory of Swine and Poultry Breeding Industry, Key Laboratory of Livestock and Poultry Multi-omics, Ministry of Agriculture and Rural Affairs, and Farm Animal Genetic Resources Exploration and Innovation Key Laboratory of Sichuan Province, College of Animal Science and Technology, Sichuan Agricultural University, P. R. China

**Keywords:** RIC-seq, transcription factors, spatial interaction, genome architecture, RNA-RNA interactions

## Abstract

We applied the latest RIC-seq technology to study the RNA-RNA interactions in human liver HepG2 cells. Integrating ChIP-seq data with the RIC-seq data, we classified vari ous transcription factors (TFs) into two distinct clusters based on the interaction enrichment scores of the uaRNA (RNA transcribed in promoter regions) and eRNA (RNA tran scribed in enhancer regions) potentially transcribed by the TF binding sites. We went on to compare the genomic features and clinical significance across different clusters, by further integrating Hi-C, ATAC-seq, and clinical data from liver cancer patients. Our results showed that Cluster 1 exhibited low spatial interaction (RIC-seq derived RNA-RNA interactions between regulatory RNA elements) and high tissue specificity (high proportion of binding sites covered by shared peaks between two different cell lines for each histone modification marker), showed higher associations with the B compartments and nucleoli, contained less densely clustered TF binding sites, and is more correlated with clinical outcomes. In contrast, Cluster 2 has higher spatial interaction, is more likely to be found in the A compartments and nuclear speckles, contained more densely clustered TF binding sites, and is more prone to form loops. These findings suggested that these transcription factors might play critical roles in regulating gene expression and serve as potential biomarkers and therapeutic targets with significant implications for cancer prognosis and treatment. In summary, groups of TFs could occur as distinct sub-networks based on spatial proximity of non-coding RNAs transcribed by the TF binding sites. These sub-networks appear to be related to regulatory differences for the TF groups and may partially explain the chromosome compartments/domains.

## Background

Transcription factors (TFs), which bind specific DNA sequences to regulate gene expression, are fundamental to development, differentiation, and the establishment of gene expression networks^[1–3]^. Some TFs exhibit versatile functions, regulating distinct genes across various cell types^[4]^, others display a high degree of specificity, acting predominantly within particular cell types or under stringent sequence- and binding-context constraints. These constraints are defined by high-throughput SELEX and ChIP-seq analyses and include motif composition, flanking nucleotide preferences, as well as requirements for dimer orientation and spacing^[5]^. Chromatin immunoprecipitation followed by sequencing (ChIP-seq) has emerged as a powerful epigenomic technique to identify TF binding sites across the genome in diverse cell types^[6, 7]^.

The regulation of gene expression by transcription factors is influenced not only by the linear distance along the DNA but also by the three-dimensional structure of the genome^[8]^. Recently developed techniques for mapping chromatin’s three-dimensional conformation include microscopic imaging and a range of capture-based approaches such as 3C, 4C, 5C, and Hi-C ^[9–13]^. These methods reveal spatial chromatin interactions, such as chromatin loops and enhancer-promoter contacts, by detecting DNA-DNA interactions^[14, 15]^. Previous studies suggest that specific transcription factors often stabilize chromatin loops by binding to their anchors, thereby regulating long-range gene regulatory networks^[16]^.

Recent studies have revealed that many active enhancer regions transcribe a class of non-coding RNAs (ncRNAs) known as enhancer RNAs (eRNAs), whose expression is strongly associated with the activation of target genes^[17–20]^. eRNAs are thought to function as mediators in the formation of chromatin loops between enhancers and promoters, thereby enhancing gene expression^[21–24]^. Additionally, eRNAs may contribute to epigenetic modifications, for instance, by regulating chromatin states through interactions with histone deacetylases (HDACs)^[25]^, which in turn modulate gene transcriptional activity^[26]^. In contrast, upstream promoter-associated RNA (uaRNA) is a class of non-coding RNA transcribed from upstream regions of gene promoters^[23]^. Functionally, uaRNAs play a role in regulating gene expression by stabilizing open chromatin structures in promoter regions, binding transcription factors, or recruiting RNA polymerase II^[27, 28]^.

In recent years, several methods for detecting RNA-RNA interactions have been rapidly developed. CLASH, by ligating the two ends of double-stranded RNA molecules, faces challenges in precisely identifying the specific regions involved in RNA duplex interactions^[29]^. HiCLIP improves upon CLASH by introducing an additional adapter, addressing some of its limitations, but still remaining susceptible to false positive results ^[30]^. MARIO captures RNA-RNA interactions mediated by RNA-binding proteins (RBPs) within cells, and may also lead to the introduction of physiologically irrelevant RNA interactions ^[31]^.

Addressing these limitations and filling the preceding technology gaps is critical for the comprehensive profiling of RNA structures and interactions. Yuanchao Xue et al.^[32]^ recently developed RNA *in situ* conformation sequencing (RIC-seq), a technique that performs *in situ* proximal RNA ligation and marks chimeric junctions of ligated RNAs using a pCp-biotin moiety, which not only enables the enrichment of chimeric reads but also allows the accurate assignment of duplex positions. In addition, RIC-seq simultaneously detects duplexes and long-range loop-loop interactions *in situ*, which is crucial for estimating higher-order RNA structures in physiological conditions. Furthermore, the synthesis of expensive probes is not required for RIC-seq, and mapping of ncRNA targets of diverse types can be achieved. Last but not least, this method offers improved sensitivity and enhanced accuracy for detecting the structures of RNAs, and enables unbiased mapping of RNA-RNA interactions at single-nucleotide resolution, thereby facilitating the reconstruction of higher-order RNA structures, the identification of direct non-coding RNA (ncRNA) targets (including lncRNAs, snoRNAs and eRNAs), and the exploration of long-range loop-loop interactions and enhancer-promoter connectivity ^[33, 34]^.

Using RIC-seq technology, we performed high-throughput sequencing of HepG2 cells and integrated publicly available HepG2 datasets to systematically analyze eRNA-uaRNA interactions. By combining this analysis with ChIP-seq, we classified 115 transcription factors (TFs) into two distinct clusters. Integrating Hi-C data, and clinical data from liver cancer patients, we further explored the genomic characteristics of these TF clusters and examined their correlation with tumor patient prognosis, shedding light on their potential role in tumor initiation and progression.

## Results

Firstly, we seek to classify transcription factors (TFs) into distinct clusters by integrating RIC-seq and ChIP-seq data. We downloaded ChIP-seq data for human HepG2 cells from the Cistrome Database, comprising 229 samples. After data preprocessing and quality control, 115 transcription factors (TFs) were retained. By integrating these data with both newly generated data and publicly available RIC-seq data(Batch-effect comparisons showed that new and public RIC-seq datasets had similar mapping rates (≈81% vs ≈86%), chimeric read rates (≈1.1% vs ≈1.0%) (quality control in Supplementary Table 1), indicating minimal batch effects between datasets.), we analyzed the interaction enrichment scores between uaRNA (promoter transcriptional RNA) and eRNA (enhancer transcriptional RNA), which are potentially transcribed from transcription factor binding sites. Interactions spanning less than 20 kb were excluded from the analysis to ensure the accuracy of the results, and the remaining transcription factors were categorized into two distinct clusters based on pairwise clustering analysis (Fig. 1a) (Supplementary Table 2).

**Fig. 1.**
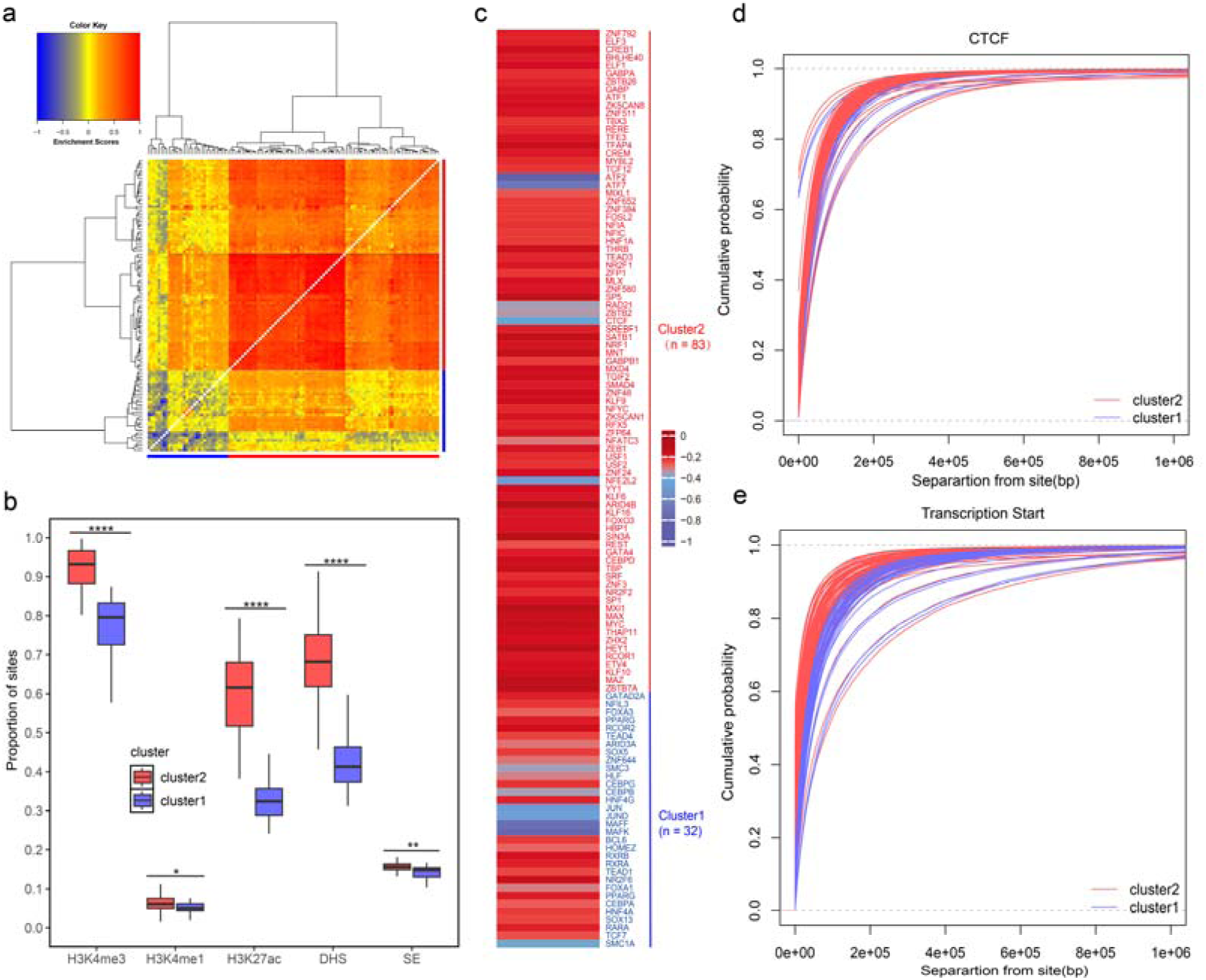
**a.** Spatial interactions between transcription factors (TFs) in the human HepG2 genome classified the TFs into distinct clusters. The normalized interaction enrichment scores between different TFs are represented as a colored matrix(TFs in Cluster 1 and Cluster 2 are indicated in blue and red, respectively.). **b.** Different conservation levels of TF epigenetic marks across the two clusters between HepG2 and hESCs.(Statistical significance was assessed using Wilcoxon rank-sum test, *P < 0.05, **P < 0.01, ***P < 0.001, ****P < 0.0001) **c.** Raw enrichment scores of the two TF clusters in chromosomal A compartment. TFs are displayed according to the hierarchical clustering order shown in panel 1a. **d.** Cumulative distribution of distances between TF binding sites and nearest CTCF binding sites in the human HepG2 genome. Each TF is represented by a separate curve, and color-coded according to its cluster. (Statistical significance was assessed using Wilcoxon rank-sum test, *P* > 0.05 for cluster1 vs cluster2) **e.** Cumulative distribution of distances between TF binding sites and nearest TSSs in the human HepG2 genome. Each TF is represented by a separate curve, and color-coded according to its cluster. (Statistical significance was assessed using Wilcoxon rank-sum test, *P* < 0.001 for cluster1 vs cluster2)

Cluster 1 comprises 32 transcription factors, which display lower degree of spatial interaction. The spatial interactions formed by these TFs suggest that their binding on the chromosome often facilitates interactions with distal regions. Among them, HNF4A and HNF4G, homologous nuclear receptor transcription factors, bind to spatially closely positioned sequences. These factors are essential regulators of key metabolic genes in the liver and pancreas.As core transcription factors specifically expressed in the liver, they play critical roles in lipid metabolism and glucose homeostasis. Studies have revealed that the chromatin regions bound by HNF4A and HNF4G are often highly overlapping, suggesting potential synergistic regulation of gene networks associated with liver function and metabolism^[35]^.

Cluster 2 consists of 83 transcription factors that exhibit relatively high interaction enrichment scores (Fig. 1a). Notably, USF1 and USF2 bind to spatially closely positioned sequences, which are upstream transcription factors from the basic helix-loop-helix (bHLH) family, typically co-regulate promoter regions as dimers and share highly overlapping binding sites^[36]^. Similarly, GABPA and GABPB1, key subunits of the GA-binding protein complex, bind to spatially closely positioned sequences and frequently associate to bind adjacent DNA elements as a complex. This interaction forms stable transcription factor networks that collaboratively regulate protein synthesis and oxidative metabolism^[37]^.

Additionally, we compared HepG2 cells with human embryonic stem cells (hESC) in terms of H3K4me3, H3K4me1, H3K27ac histone markers, DNase I hypersensitive sites (DHS), and super-enhancers (Fig. 1b). The overall trend observed was TF in Cluster 2 > Cluster 1 (Figure 1b), indicating that transcription factors in Cluster 2 exhibit a stronger binding affinity within open chromatin regions conserved between cell lines.

Using Hi-C data, we examined the distribution of transcription factors (TFs) within chromosomal A and B compartments(Fig. 1c). A compartments are generally associated with euchromatin, characterized by active transcription, while B compartments are more heterochromatic and contain genes with lower transcriptional activity. We validated the accuracy of A/B compartment assignment using gene density and histone modifications (H3K27ac) (see Supplementary Fig. 1). We calculated the proportion of the binding sites for each TF located within these active regions. The results indicate that Cluster 1 TFs are more frequently distributed in B compartments, aligning with their association with less transcriptionally active regions. This observation is consistent with findings reported by Xiaoyan Ma et al.^[38]^. These factors participate in critical cellular processes, including cell proliferation and apoptosis. In contrast, Cluster 2 TFs are relatively more enriched in A compartments (*P* < 0.0001 for Cluster 2 vs Cluster 1), which are highly transcriptionally active and closely linked to actively transcribed genes.

Gene Ontology (GO) analysis revealed that transcription factors in Cluster 1 are predominantly involved in liver development and metabolic regulation. Several key biological pathways were significantly enriched, including the nuclear receptor transcription pathway (*P* = 8.56×10□¹□) and adipogenesis (*P* = 6.40×10□¹□), both of which play central roles in hepatic metabolism, cell proliferation, and tumorigenesis. In addition, the p53 signaling pathway (*P* = 1.15×10□□), a critical tumor suppressor pathway in liver cancer development, serves an important protective function.In contrast, GO analysis of Cluster 2 indicated that transcription factors are mainly implicated in fundamental transcriptional regulatory processes, including regulation of miRNA transcription (*P* = 4.72×10), DNA-templated transcription (*P* = 6.99×10□¹²), and nerve growth factor (NGF)-stimulated transcription (*P* = 1.08×10□□). These results suggest that TFs in Cluster 2 primarily function as core regulators of basal transcriptional programs.

We identified target genes of TFs using BETA^[39]^ and performed functional enrichment analyses for Cluster 1 and Cluster 2. Cluster 1 targets were mainly involved in hepatocyte metabolism and lipid homeostasis, including PPAR signaling(*P* = 6.13×10^−10^), lipoprotein transport(*P* = 2.9×10^−11^). Cluster 2 targets were enriched in stress response(*P* = 3.4×10^−6^), apoptosis and necroptosis(*P* = 2.8×10^−10^).

The two clusters of transcription factors (TFs) exhibit distinct linear relationships with respect to their chromosomal locations and distances from specific gene regulatory elements. Namely, transcription factors in Cluster 2 shows a tendency toward shorter distances to CTCF binding sites (Fig. 1d), whereas they were significantly closer to transcription start sites (TSSs) (Fig. 1e). This proximity suggests that they may directly regulate gene promoter regions. In contrast, Cluster 1 TFs are located farther from these key regulatory elements, implying a more indirect role in gene activation and transcriptional regulation. The greater distances observed for Cluster 1 TFs also align with their lower spatial interaction requirements.

In summary, transcription factors in Cluster 2 play a more active role in regulating gene transcription and exhibit stronger spatial interactions within regions of active transcription. These findings highlight the functional distinctions among the clusters in their contributions to chromosomal organization and transcriptional regulation.

We went on to examine various genomic features for distinct clusters. Through the analysis of histone H3K27ac, we identified the peak regions of super-enhancers and quantified the enrichment levels of transcription factors (TFs) in super-enhancers. The results revealed that TFs in Cluster 1 exhibited higher overlap levels within super-enhancer regions (*P* < 0.001 for cluster1 vs cluster2), conversely, Cluster 2 demonstrated lower enrichment, suggesting that super-enhancers may play a more critical role in transcriptional activation for TFs in Cluster 1(Fig. 2b).

**Figure 2a.**
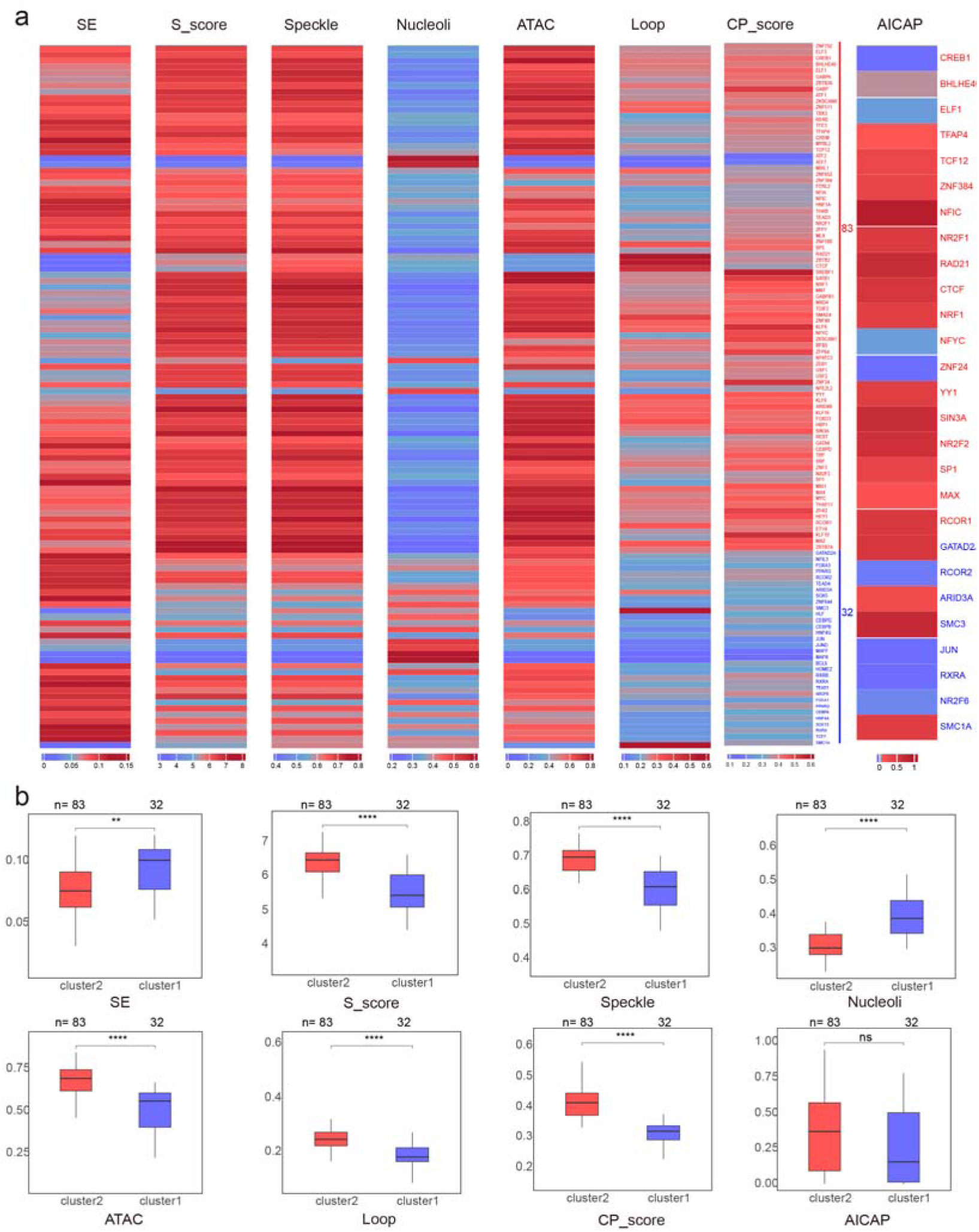
This figure illustrates the enrichment patterns of super-enhancers(SE), S scores, speckles, nucleoli, ATAC peaks(ATAC), Hi-C loop anchors(Loop), CP scores and AICAP scores across different clusters. SE shows proportion of TF binding sites covered by super-enhancers, S scores were average of S_basis scores estimated using HiCAN for each TF (with higher values indicating higher propensity to be in Speckle phase), speckles were proportions of TF binding sites covered by the Speckle-associated regions, nucleoli were proportions of TF binding sites covered by the Nucleolus-associated regions, ATAC shows proportion of TF binding sites covered by ATAC-seq peaks, Hi-C loop anchors were proportions of TF binding sites covered by the Hi-C loop anchor regions, CP scores were average of clustering propensity scores estimated using previously described method^[48]^ for each TF, AICAP scores were average of anti-1,6-HD index of of chromatin-associated proteins (AICAP) indicating phase-separation potential directly downloaded from public resources^[49, 50]^. Here raw values were all normalized to N(0,1) distribution by subtracting by average and divide by standard deviation for each column. **Figure 2b** This figure depicts the distribution of super-enhancers, S scores, speckles, nucleoli, ATAC peaks, Hi-C loop anchors, CP scores and AICAP scores across various clusters. Here raw values were shown and subjected to statistical tests. Different clusters are marked with distinct colors. Inter-group differences were assessed using the Wilcoxon rank-sum test, with significance levels denoted by asterisks (*P < 0.05, **P < 0.01, ***P < 0.001, ****P < 0.0001).

ATAC-seq, a technique used to analyze the genome by detecting open chromatin regions, provided further insights into chromatin accessibility^[40]^. Our findings showed that TFs in Cluster 2 exhibited higher enrichment in ATAC-seq peaks (*P* < 0.0001 for cluster2 vs cluster1), indicating that TFs in this cluster are more likely to bind within open chromatin regions. These regions are characterized by high transcriptional potential and are strongly associated with gene expression activity. In contrast, Cluster 1 displayed lower chromatin accessibility, suggesting that the TFs in this cluster are prone to bind heterchromatin regions with specialized functions.

Analysis of nucleolus and speckle localization revealed that transcription factors (TFs) in both Cluster 2 predominantly reside in speckles. TFs in the nucleolus are often associated with non-coding RNA processing and ribosome synthesis, while nuclear speckles are regions enriched in transcriptional and splicing activity^[41]^. These findings suggest that TFs in Cluster 2 may play a more important role in transcriptional activation and splicing regulation.

In Cluster 1, two TFs, MAFF and MAFK, exhibit over 50% binding site overlap with nucleolar localization. These TFs are implicated in nucleolus-relate functions, including ribosome biogenesis and the regulation of the cellular stress response^[42]^. Moreover, MAFF and MAFK are likely involved in the formation of homodimers; however, due to their lack of transactivation domains, they may function as transcriptional repressors. This inhibition of transcriptional activity aligns with their roles in transcriptional regulation within the nucleolus^[43]^.

Analysis of chromatin loops using Hi-C data revealed that CTCF, RAD21 in Cluster 2 exhibit higher propensity for loop formation, with Cluster 1 showing lower tendency (*P* < 0.0001 for cluster1 vs cluster2). CTCF and RAD21 are well-known components of chromatin loop structures. These factors are integral to the spatial organization of transcriptional machinery, facilitating chromatin looping that is often closely associated with gene expression regulation and cell division. As key members of the chromatin loop complex, RAD21 contribute to the regulation of loop stability, thereby influencing chromosome structure and function^[44]^. In contrast, loop formation potential for MAFF and MAFK in Cluster 1 are low, suggesting their involvement in transcription repression.

The CPscore is a quantitative measure of transcription factor (TF) clustering propensity of their binding sites. Analysis of CPscore revealed that Cluster 2 exhibited relatively high scores (*P* < 0.0001 for cluster2 vs cluster1), whereas MAFF and MAFK in Cluster 1 showed lower scores. The consistency between chromatin’s three-dimensional interactions and its linear structure underscores the dual regulation of TF activity by forming clusters based on: (1) linear proximity and (2) spatial vicinity. These findings highlight their potential role as critical regulators of gene expression in promoter and enhancer regions^[45]^.

In the analysis of the anti-1,6-HD index (AICAP) for chromatin-associated proteins, Cluster 2 displayed higher values (*P* > 0.05 for cluster2 vs cluster1), with Cluster 1 showing lower values. Notably, CTCF and RAD21 are recognized as key factors involved in chromatin structure and phase separation^[46]^. By interacting with other proteins, these factors contribute to the formation of physical domains, regulate the three-dimensional organization of chromatin, and promote chromatin phase separation^[47]^.

In conjunction with clinical data from patients with liver cancer, we conducted a predictive analysis of eRNA expression across different clusters. The analysis indicated that cluster 1 exhibited higher proportions of eRNAs whose expression was associated with clinical outcomes (*P* < 0.0001 for cluster1 vs cluster2), and this enrichment suggests that eRNAs associated with Cluster 1 may have stronger associations with hepatocellular-related regulatory programs. In contrast, cluster 2 exhibited lower proportion of predictive eRNAs.

Interestingly, the expression levels of MAFF (survival *P* = 0.0015) and MAFK (survival *P* = 0.0466) itself in cluster 1 were also significantly associated with clinical outcomes in liver cancer (see Supplementary Fig. 2). Notably, MAFF may contribute to tumor-initiating cells (TICs), whose functions are critical in determining tumor aggressiveness and metastatic potential^[51]^. MAFF expression levels showed an association with patient prognosis, suggesting that MAFF may serve as a potential prognostic candidate in liver cancer. Moreover, in the tumor microenvironment of liver cancer, hypoxia frequently induces MAFF expression through activation of the HIF pathway^[52]^. MAFF subsequently drives transcriptional programs associated with tumor malignancy by activating the IL11 and STAT3 signaling pathways ^[53]^.

We further applied BETA to predict putative target genes of MAFF and MAFK. Functional enrichment analysis revealed that MAFF putative target genes were significantly enriched in the NRF2 pathway(*P* = 2.53×10^−7^), ferroptosis(*P* = 1.32×10^−4^). MAFK target genes were preferentially enriched in NF-kB activation(*P* = 4.7×10^−4^), apoptosis(*P* = 2.1×10^−4^).

Based on the above analysis, we selected three MAFF-binding peak regions with eRNA transcription and three putative target genes regulated by MAFF for in-depth investigation (Fig. 3). These target genes have been reported to participate in cancer-related signaling pathways. SNX5 belongs to the sorting nexin family and is primarily responsible for endosomal membrane trafficking and signal transduction. Studies have demonstrated that SNX5 is aberrantly expressed in various types of cancer and may influence cancer cell proliferation and invasion by regulating intracellular protein transport pathways. For example, SNX5 may influence cancer cell migration and proliferation by modulating the EGFR (epidermal growth factor receptor) signaling pathway^[54]^. RNF24 is a membrane protein that interacts with TRPC (transient receptor potential channel) proteins^[55]^. In certain cancers, RNF24 has been identified as an oncogenic factor, particularly in esophageal cancer and oral squamous cell carcinoma^[56]^. RAB10, a member of the Ras superfamily, is primarily involved in intracellular vesicle transport^[57]^. Research indicates that RAB10 contributes to the progression of various malignancies, including liver cancer, cervical cancer, and gliomas, through mechanisms such as acting as a target for non-coding RNAs, regulating the AMPK (AMP-activated protein kinase) signaling pathway, and modulating autophagy^[58].^

**Figure 3.**
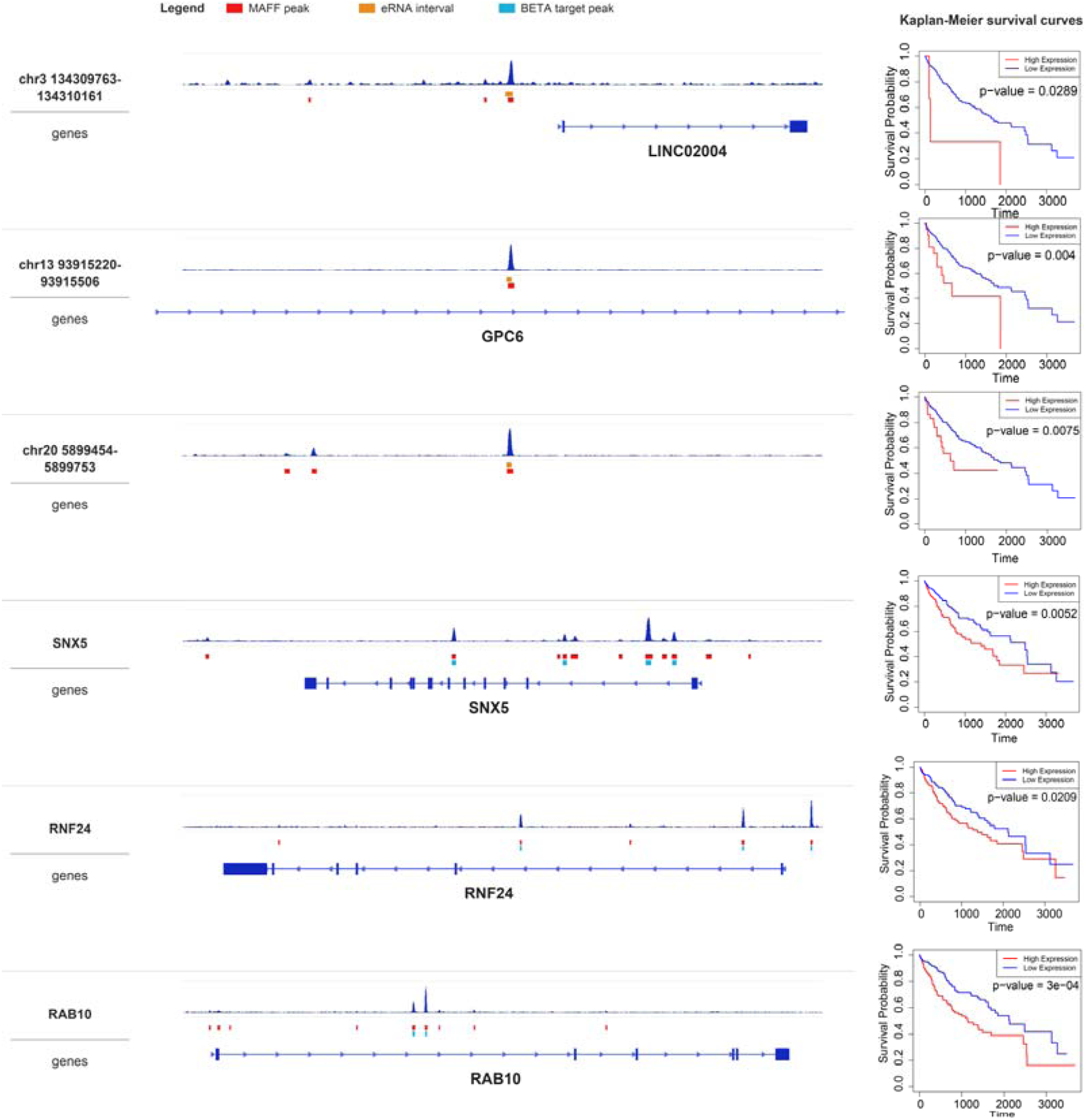
**The left panel** illustrates three selected eRNA peak regions of MAFF along with their three target genes. These visualizations highlight peak intensity and intervals, as well as gene annotation and eRNA peak regions. The orange boxes represent eRNA loci, the red boxes indicate MAFF ChIP-seq peak regions, and the blue boxes denote target gene peak regions **The right panel** presents the survival analysis results for the aforementioned eRNAs and genes, with statistical significance (*P-*values) assessed using the log-rank test. The survival curves provide a clear comparison of survival outcomes between highly expressed and lowly expressed groups. The *P*-value underscores the statistical significance of the observed differences in survival between the two groups.

In this study, we utilized the NetworkX package in Python to further investigate the regulatory network and identify hub genes. The top 5% of genes with the highest degree were selected as hub genes, and eRNAs and uaRNAs were extracted. Subsequently, we visualized the top seven hub-uaRNAs with the highest number of interactions and all the identified hub-eRNAs. Additionally, we explored the overlap between these genes and transcription factor (TF) binding sites. For each cluster, the proportions of hub-eRNAs and hub-uaRNAs were calculated and visualized using boxplots (Fig. 4). The Wilcoxon rank-sum test revealed significant differences in overlap proportion between clusters. In hub-uaRNAs and hub-eRNAs, the difference between Cluster 1 and Cluster 2 was statistically significant (*P* < 0.05).

**Figure 4.**
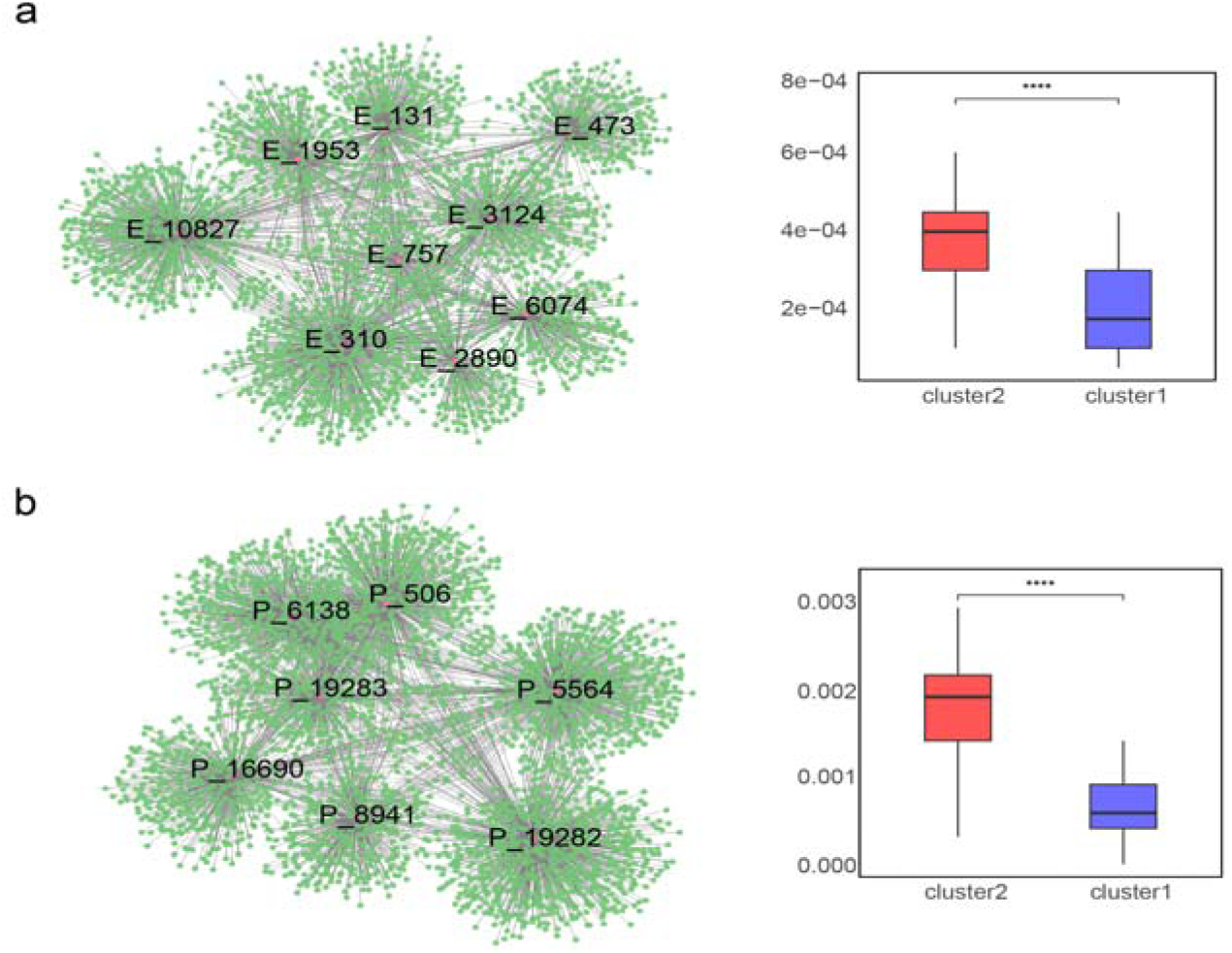
The network diagram on the left illustrates the connectivity degree of hub genes (hub-eRNAs and hub-uaRNAs) (**4a and 4b**), whereas the boxplots on the right depict the overlapping degrees (proportions of TF binding sites covered by hub-eRNA/hub-uaRNA for each TF) between hub-eRNA/hub-uaRNA (**4a and 4b**) and TF binding sites across the two distinct clusters(Statistical significance was assessed using Wilcoxon rank-sum test, *P < 0.05, **P < 0.01, ***P < 0.001, ****P < 0.0001)

Furthermore, we selected the top three eRNAs and uaRNAs with the highest highest degrees and performed Gene Ontology (GO) analysis on their interacting mRNAs. The results indicated that eRNAs may influence gene expression by regulating ribonucleoprotein complex biogenesis(*P* = 3.16×10^-6^), protein degradation(*P* = 3.16×10^-5^), and mRNA metabolic pathways(*P* = 1.26×10^-4^). In addition, eRNAs are also associated with intracellular protein transport(*P* = 1.26×10^-5^) and cell division(*P* = 3.16×10^-4^), potentially modulating molecular signaling through vesicle and membrane transport to regulate the cell cycle. In contrast, uaRNAs are implicated in critical biological processes such as chromatin remodeling(*P* = 2.08×10^-6^), cell cycle regulation(*P* = 3.70×10^-4^), and DNA damage response(*P* = 6.21×10^-7^). By modulating chromatin structure and gene expression, uaRNAs appear to play pivotal roles in cell cycle progression, contributing to cell proliferation, division, and genomic stability. These functions are closely linked to tumor development and progression.

## Discussion

The latest RIC-seq technique enables the capture of protein-mediated RNA-RNA proximal interactions in living cells at single-base resolution. By analyzing pairs of interacting enhancer RNAs (eRNAs) and promoter-derived non-coding RNAs (uaRNAs), this approach facilitates the identification of enhancer-promoter interactions. Integration with ChIP-seq and Hi-C data has revealed that transcription factor (TF) binding sites are not only governed by DNA sequence specificity but are also intricately associated with the three-dimensional organization of chromatin. Notably, eRNAs exhibit synergistic regulatory functions in facilitating the formation of chromatin loops between enhancers and promoters, underscoring their critical role in chromatin architecture and gene regulation.

We employed RIC-seq technology to perform high-throughput sequencing of HepG2 cells and integrated publicly available HepG2 data sets to systematically analyze intermolecular interactions. Additionally, by incorporating ChIP-seq, Hi-C, ATAC-seq, and clinical data of patients with liver cancer, we classified 115 transcription factors (TFs) into two distinct clusters. Cluster 1 exhibited low spatial interactions and low cell conservation (proportions of binding sites covered by shared peaks between 2 different cell lines (hepG2 and ESC) of each histone modification marker), as well as higher association with B compartment and nucleoli. This cluster featured a lower density of TF binding sites and demonstrated stronger correlations with clinical outcomes. In contrast, Cluster 2 displayed higher spatial interactions, was preferentially associated with A compartments and nuclear speckles, and contained a higher density of TF binding sites.

The varying overlap proportions of binding sites of different transcription factors (TFs) with these two compartments reflect fundamental differences in TF function, target gene regulation, transcriptional dynamics, and cellular homeostasis. TFs with higher association with speckles are primarily involved in protein-coding gene expression, co-transcriptional processing, and alternative splicing. For example, binding sites of splicing-coupled TFs overlap with speckles to co-localize with splicing machinery^[59]^. This ensures that pre-mRNA splicing is coordinated with transcription elongation-critical for maintaining transcript fidelity and isoform diversity. Elongation-support TFs overlap with speckles and are involved in productive elongation^[60]^. TFs with higher association with nucleoli specialize in ribosome biogenesis and Pol I-mediated transcription (the rate-limiting step for ribosome production)^[61]^. Speckles act as “hubs” for rapid mobilization of splicing and transcription factors^[62]^. TFs with high speckle overlap can quickly recruit co-factors to target promoters/enhancers, enabling rapid, stimulus-dependent induction of protein-coding genes (e.g., immune response genes, heat shock proteins). Nucleolus-associated TFs: Nucleoli are transcriptionally constrained compartments (only Pol I activity is dominant). TFs with high nucleolar overlap are restricted to regulating rDNA, avoiding crosstalk with Pol II-mediated transcription^[61]^. TFs that bind euchromatin (active histone marks: H3K4me3, H3K27ac) have higher speckle overlap^[63]^. In contrast, nucleoli are flanked by heterochromatin (NOR-associated heterochromatin)^[64]^.

These findings align with the results reported by Xiaoyan Ma et al.^[38]^, providing further validation and support for the spatial behavior and distribution of TF binding in chromatin. We argued that new biological insight is gained beyond previous Hi-C/TF interaction cluster studies, and TFs by RIC-seq-derived RNA interactions is meaningful mechanistically. Firstly, Hi-C focuses on chromatin spatial proximity (DNA-DNA interactions), and traditional TF clustering relies on TF binding sites (ChIP-seq) or protein-protein interaction-these approaches have inherent limitations that the eRNA/uaRNA + TF clustering integration overcomes. The integration of eRNA/uaRNA pairing with TF clustering moves beyond static DNA topology (Hi-C) to reveal dynamic, RNA-dependent functional TF modules that drive gene expression-resolving the gap between TF binding (observed by Hi-C/ChIP) and TF function (validated by RNA interactions). Secondly, eRNAs possess intrinsic structural features that enable them to serve as scaffolds for recruiting multiple transcription factors^[65]^. RIC-seq/ChIP-seq identifies RNA regions that physically interact with TFs; clustering TFs bound to the same RNA scaffold reveals a “TF-RNA complex” where the RNA stabilizes TF-TF interactions (e.g., TF A and TF B bind interacting eRNAs from the same RNA scaffold, and the RNA bridges their physical interaction). This clustering is not arbitrary -it reflects a biochemically stable TF complex assembled on an RNA scaffold, which directly drives transcriptional activation/repression. Clustering transcription factors based on RIC-seq RNA interactions is meaningful, as it highlights the role of RNA as a scaffold for the assembly of physiologically relevant TF complexes. Thirdly, dysregulation of eRNA/uaRNA is a hallmark of diseases (e.g., cancer, neurodegeneration). Clustering TFs by RNA interactions identifies disease-specific TF modules driven by ncRNA. For example, in breast cancer, an oncogenic eRNA pairs with uaRNA at the MYC locus, and clustering TFs bound to this RNA reveals a TF module (e.g., c-Myc, E2F1, FOXM1) that is up-regulated in tumors. Targeting the eRNA disrupts the TF cluster and reduces MYC expression -proving mechanistic causality. This clustering is meaningful, as it identifies actionable TF modules (and their RNA scaffolds) for therapeutic intervention-capabilities that traditional TF clustering approaches cannot provide.. Our approach uncovers a new layer of gene regulation -- RNA-dependent TF cluster plasticity -- that links ncRNA dysregulation to TF network rewiring in disease, a discovery not possible with prior methods.

Although this study elucidates the spatial distribution characteristics of transcription factors in HepG2 cells and their potential association with liver cancer prognosis, several limitations should be acknowledged. Firstly, our conclusion is based solely on computational results, and further experimental validations are required to confirm our biological claims. The necessary experiments include but not limited to validation of selected eRNA-uaRNA interactions, functional validation of MAFF-regulated eRNAs, protein binding or TF dependency experiments. Secondly, Only one cell line (HepG2) was used in our study and lack of comparative data (e.g., normal cells or other cancer cell lines) limits the generalizability of our findings. Admittedly tumor has heterogeneity in both spatial zonation and cell type composition, but hepG2 is a clonal line derived from a single individual’s liver tumor, with limited heterogeneity in aspects of genetics, epigenetics, phenotypes and functions, which are likely the results of subtle difference in culture conditions and experimental manipulations between distinct labs where this cell line was maintained. HepG2 is commonly used in vitro model of human hepatocytes with limited heterogeneity. However, the intrinsic heterogeneity of HepG2 cells should be acknowledged, and the conclusions of our study could be strengthened by future study using single-cell level multi-omics technologies. Thirdly, our study is based on computational enrichment analyses of correlative factors, is limited by annotation dependence and lacks cell-type specificity. Fourthly, GO analysis on a large and heterogeneous list of transcription factors can yield broad, non-specific enrichments that reflect generic transcription factor functions rather than meaningful, cluster-specific biology. In summary, future studies should incorporate more extensive experimental validation and comparative analyses across different cell types to strengthen the reliability and generalizability of these conclusions.

## Methods

### Cell culture

HepG2 cells (ATCC, HB-8065) were cultured at 37 °C in a humidified incubator with 5% CO. Cells were maintained in DMEM supplemented with 10% fetal bovine serum and 100 U ml ¹ penicillin–streptomycin. Cells were confirmed to be mycoplasma-free by PCR-based detection.

### Promoter-enhancer interaction(PEI) identification

The ChIP–seq data of H3K4me3 and H3K27ac for hepG2 cell line were downloaded from the ENCODE^[66]^. The data were processed using Bowtie2(v2.4.4)^[67]^ and the resulting SAM files were then sorted with Samtools(v1.6)^[68]^. Peak calling was performed using MACS2^[69]^ software. Super-enhancers were defined based on the coverage of H3K27ac ChIP-seq signals using the software Rank Ordering of Super-Enhancers (ROSE)^[70]^.

We utilized both self-generated data and publicly available RIC-seq datasets of hepG2 cell line and carried out analyses to identify inter-molecular RNA-RNA interactions as previously described^[34]^. In brief, RIC-seq libraries were generated following the protocol described by Yuanchao Xue et al^[34]^. And adapter sequences were removed using the Trimmomatic program (v0.36)^[71]^. Secondly, PCR duplicates were removed using custom scripts, and low-complexity fragments were trimmed from the ends of reads using Cutadapt (v1.15)^[72]^. Thirdly, reads were mapped to rRNA sequences using STAR software (v2.7.10a)^[73]^. Fourthly, we constructed an index of the human reference genome and enhancer-promoter sequences. The remaining reads were aligned to the human reference genome using the STAR software, obtaining normally mapped reads (Aligned_out.sam) and chiastically mapped reads (Chimeric_out.sam). Intermolecular sequences were subsequently identified and extracted. Ultimately, high-confidence intermolecular RNA-RNA interactions were obtained.

### Identification of transcription factor binding sites

The ChIP-seq narrowPeak data sets for the transcription factors (TFs) in HepG2 cells were obtained from the Cistrome database^[74]^. We focused on autosomal and sex chromosomes X and ranked the peaks based on their scores (minus log transformed FDR values) in descending order. The top 20,000 regions were selected for subsequent analyses.

### Clustering analysis

We classified various transcription factors (TFs) into distinct clusters based on the interaction enrichment scores of uaRNA (RNAs transcribed in promoter regions) and eRNA (RNAs transcribed in enhancer regions) potentially regulated by TF binding sites. Firstly, we extracted three types of interactions-PPI (Promoter-Promoter Interactions), PEI (Promoter-Enhancer Interactions), and EEI (Enhancer-Enhancer Interactions)-from the RIC-seq results. Interactions with distances over 20 kb were retained.

We performed enrichment analysis based on the binding sites of transcription factors (TFs) and the RNA-RNA interactions (PPIs, PEIs and EEIs). A custom script (genomic regions were considered overlapping when the coordinates of two regions shared at least one base pair overlap)was used to calculate the number of RNA-RNA interactions between eRNA-uaRNA/eRNA-eRNA/uaRNA-uaRNA pairs with one end bond by one TF and the other end bond by the other TF (deemed as result1 here). We then grouped 300bp genomic regions with sliding step 20 bp into 5 quantiles based on GC content and mappability, respectively. Similarly, we further grouped 300bp genomic regions with sliding step 20 bp into 2 groups based on chromatin accessibility (accessible or not accessible) and A/B compartment status (A or B compartment). We then re-sized all binding sites of TFs to 300-bp length by extending upstream 150 bp and downstream 150 bp from the peak summit. For a pair of TF A and TF B, we first fixed TF A and generated shuffled TF B background sites to calculate ‘shuffle1’ value. In brief, by custom coding, we generated TF B background sites from sampling the same number of genomic sites as TF B, and the sampled binding sites also followed exact same distribution of genomic covariates as TF B, that is, the number of background sites falling in each group of GC content, mappability, chromatin accessibility and A/B compartment status should be the same as that for TF B. The number of RNA-RNA interactions between eRNA-uaRNA/eRNA-eRNA/uaRNA-uaRNA pairs with one end bond by TF A and the other end bond by the background of TF B was estimated (deemed as shuffle1 here). Similarly, fixed TF B and generated shuffled TF A background sites to calculate ‘shuffle2’ value. The average of these two values ‘shuffle1’ and ‘shuffle2’ was calculated, and the enrichment score for this TF pair was computed using the formula:

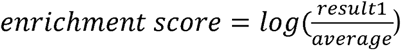

We performed shuffling 100 times each for background 1 and background 2. P-values were estimated based shuffling results, that is, the fraction of shuffles with higher supported RNA-RNA interaction number than the ‘result1’ value, and the P-values were estimated for background 1 and background 2, respectively, and the larger of the 2 P-values was used as the final P-value. The process was then repeated for every pair of TF data sets, and the pair-wise enrichment scores were transformed into a matrix using a custom script. Before clustering, enrichment scores were normalized to the range of −1 to 1. The diagonal elements corresponding to TF self-interactions were set as missing values (NA). Ultimately, hierarchical clustering based on euclidean distances and Ward.D2 linkage was applied to classified various transcription factors (TFs) into two distinct clusters. FDRs were obtained by correcting P-values for pair-wises comparisons.

### A/B compartment identification

To analyze the compartmentalization of genomic regions, we first pre-processed the bedGraph file (retrieved from 4Dgenome database) by removing NaN values from the data. Subsequently, we classified genomic regions into two compartments: compartment A and B with positive and negative indexes. To identify contiguous regions within each compartment, we use the following command to merge adjacent regions: bedtools merge -i BED. After determining the partitioning of the A and B compartments, we proceeded by calculating the overlap between TF binding sites and A compartment (Aoverlap), as well as that between TF binding sites and the A+B compartments (ABoverlap). The final enrichment score is then computed using log(Aoverlap / ABoverlap).

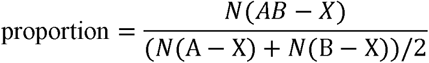

### Identification of histone mark peaks

We retrieved ChIP-seq data sets of histone modifications and open chromatin regions for HepG2 and hESC cells from the Cistrome database, including H3K4me3, H3K4me1, H3K27ac and DNase I hypersensitive sites (DHS). Super-enhancers were obtained from H3K27ac data set using the preceding method (see **PEI identification** for details). Initially, overlap intervals between a specific modification from cell type A TF X were calculated (A-X file). These two files from cell type A (A-X) and B (B-X) were subsequently overlapped to generate a final result file (AB-X). Finally, the overall enrichment value was calculated by dividing the number of peaks in AB-X by the average of that in A-X and B-X.

## Distance distribution

Transcription start sites (TSSs) were extracted from the gtf file downloaded from Ensemble 110 (https://ftp.ensembl.org/pub/release-110/gtf/homo_sapiens/Homo_sapiens.GRCh38.1 10.gtf.gz), and the nearest distance between every peak region of each transcription factor (TF) and the TSS was calculated to obtain the distance distribution of each TF relative to the TSSs. The distance distribution of each TF relative to the CTCF binding sites was similarly estimated. To visually present these distance distribution data, we used the cdf package in R(v4.4.1) to generate cumulative distribution function (CDF) plots.

### Speckle/ Nucleoli identification

The interaction matrix for each pair of chromosomes (i, j) was generated the from Hi-C file using the Juicer^[75]^ command. Missing values and columns/rows in the matrix were then filled using a custom script, and the matrix was then transposed. Finally, a large genome-wide matrix was generated from the pair-wise interaction matrices according to chromosome orders. Based on this large matrix, Hi-C interchromosomal contact map analysis was performed with NMF (GitHub: kaistcbfg/HiCAN), which identifies three gene density regions: S (Speckle-associated), N (Nucleolus-associated), and U (Undefined). In brief, the core design principle of HiCAN is mainly composed of three steps: construction of an intra-chromosomal interaction-filled inter-chromosomal Hi-C contact map, projection of the contact map into three low-dimensional spaces with NMF, and annotation of S (speckle-associated)-, N (nucleolus-associated)-, and U (undefined)-basis based on gene density. The proportions of TF binding sites overlapping with S and N types of regions were calculated. Finally, each region was scored for tendency to be Speckle-associated, and average of the scores for binding sites of each TF was estimated (S_basis score).

### ATAC-seq peak calling

The downloaded ATAC-seq data were aligned to the genome using Bowtie 2. Next, Samtools was used to remove alignments mapped to the mitochondrial genome. The filtered alignments was then sorted. Duplicates were removed an d MACS2 was employed to call peaks.

### Loop identification

To generate the loops, the Hi-C files were processed using Juicer HICCUPS^[75]^. The peak regions of transcription factors (TFs) were then overlapped with either end of the loops to assess their degree of enrichment.

### Clustering propensity (CP) score estimation

We calculated CP score as previously described^[48]^. In brief, distribution of log10 transformed distances between each TF binding site and its nearest site was estimated, and two-sided K-S test was applied to compare the distribution of ChIP-seq binding profile (T) and that of the control (C), and obtained the CP score. 100 random sets of genomic intervals are generated as controls (C) to obtain 100 CP scores for each TF. The average of the 100 CPs was used as the final CP score. Bedtools was used to find the closest distance. The CP score is defined as follows:

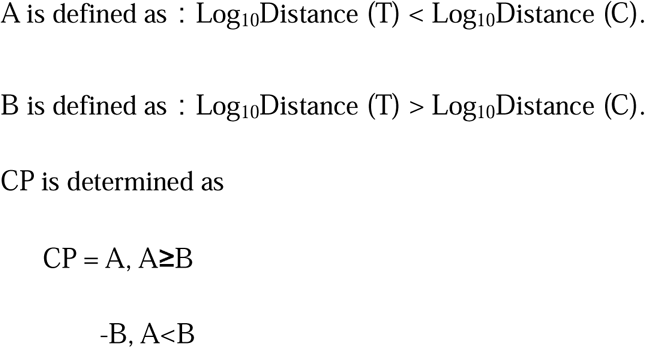

### Survival analysis

Clinical data for liver cancer patients were downloaded from a publicly available database (GDC Data Portal), which included basic patient information, tumor characteristics, and clinical follow-up data. eRNA expression data were also obtained from The Cancer eRNA Atlas (TCeA)^[76]^. By integrating eRNA expression data with clinical data, we performed a Cox regression analysis adjusting clinical covariates to assess the prognostic potential of eRNA. Age and pathological clinical characteristics, including AJCC pathological stage, T stage, N stage, and M stage, were included as covariates in the regression models.The proportion of prognostic eRNA (defined as those with *P* < 0.05) over all the annotated eRNA overlapping TF binding sites was estimated.

The minus function of BETA^[39]^ tool was then used to predict the target genes of the TFs. This tool enabled us to efficiently identify the potential target genes regulated by each TF.

Next, three prognostic eRNA peak regions and three prognostic target genes of the MAFF were visualized. Relevant location data, including the eRNA peak regions and the genomic coordinates of the target genes, were imported into IGV^[77]^, along with corresponding ChIP-seq data and genomic annotation information.

### Hub nodes analysis

We used NetworkX^[78]^ in Python to further identify the hub nodeswithin the regulatory network. Firstly, we constructed a control network based on the previously RIC-seq obtained intermolecular interaction list data. In this network, nodes represent RNA-associated elements, including gene-associated RNAs, eRNAs, and uaRNAs, and edges represent their regulatory relationships. By calculating the connectivity degree of each node within the network, nodes were ranked according to their degree values, and the top 5% nodes with the highest connectivity were selected as hub nodes. Hub eRNAs and uaRNAs were further identified from the hub nodes, and their corresponding interacting RNA partners were extracted from the original RIC-seq interaction pairs to construct hub RNA-associated regulatory networks. We used Cytoscape(v3.9.1)^[79]^ software to visually demonstrate the overall structure of the regulatory network.

## Data availability

ChIP-seq data of H3K4me3 and H3K27ac for HepG2 cells were downloaded from Cai et al^[32]^. ChIP-seq narrowPeak profiles of TFs for HepG2 cell line were retrieved from the Cistrome Data Browser (<u>Cistrome DB</u>). RIC-seq data was obtained from the public database GEO (GSE190214)^[34]^ and newly genera ted data was deposited in GEO (GSE286140). Hi-C data for hepG2 cells was downloaded from https://data.4dnucleome.org/experiment-set-replicates/4DNESC2DEQIJ/. An anti-1,6-HD index of chromatin-associated proteins (AICAP)^[49]^ were obtained from a previous study^[50]^. The ATAC-seq data were downloaded from the GEO database (GSE262479)^[80]^. Clinical data of liver cancer patients were downloaded from https://portal.gdc.cancer.gov/. eRNA expression data were obtained from https://bioinformatics.mdanderson.org/public-software/tcea/.

## Code availability

Code availability codes used for data analysis in this paper can be found at github (Yuan-68/RIC).

## Author contribution

QZ.T. and MZ.L. designed bioinformatics analyses and supervised the study. XY.L.., Y.D., YH.J., YL.T., C.W. and J.X. collected the data. Y.Y. and HD.H. performed the data analysis. QZ.T. and Y.Y. developed the analytical code and performed data visualization. QZ.T. and Y.Y. prepared the manuscript. Y.Y. revised the manuscript.

## Acknowledgements

This work was supported by the National Key R & D Program of China (2022YFF1000100 and 2020YFA0509500), the National Natural Science Foundation of China (32225046, 32421005 and 32494802), the Sichuan Science and Technology Program (2021ZDZX0008 and 2021YFYZ0009), and the Dual Support Plan for Discipline Construction—Special Program for The Cultivation of Outstanding Young Scholars (2022SZYQ004).

## Supplementary Information

**Supplementary Figure 1.**
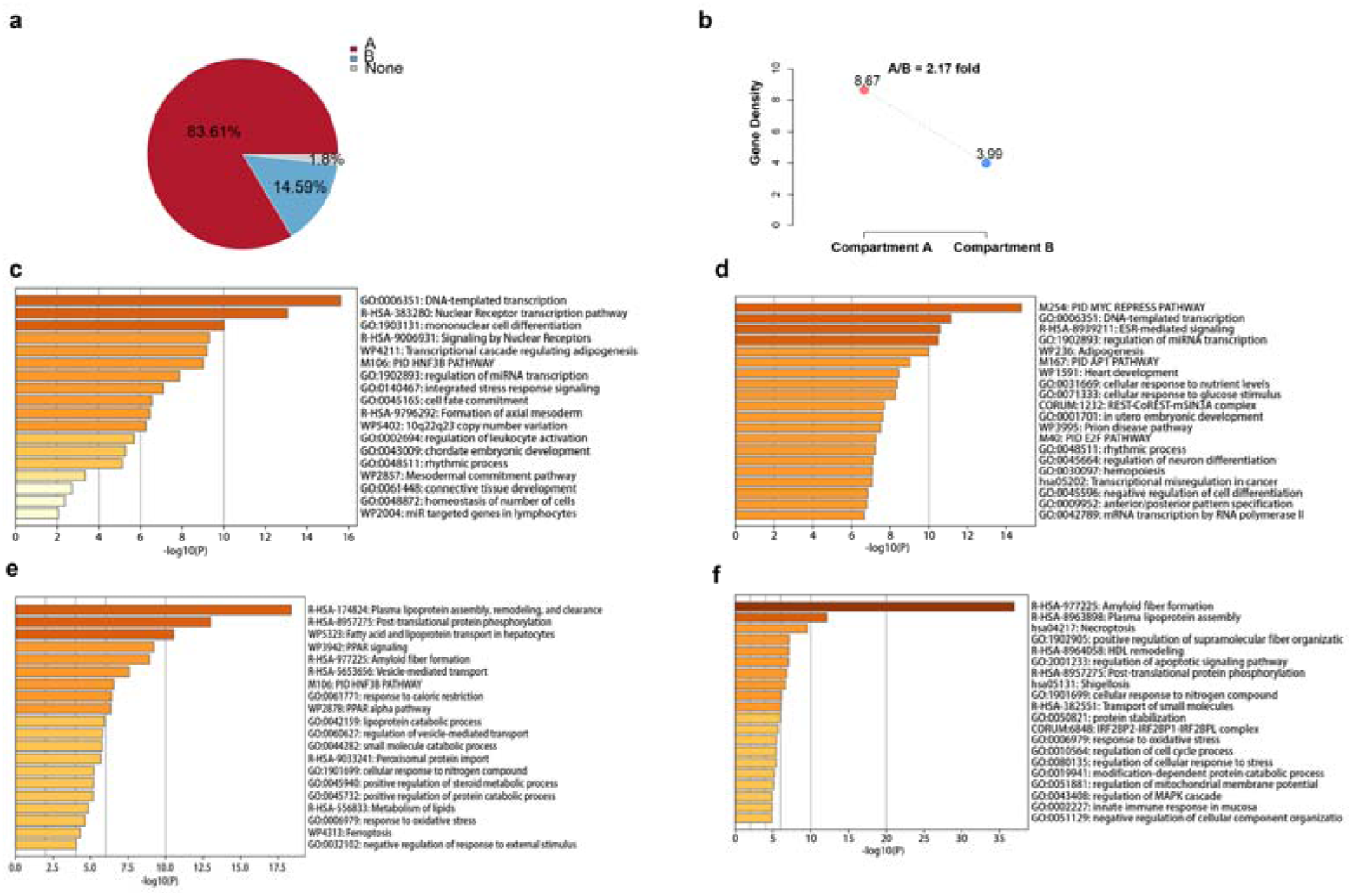
a. The proportion of the A and B compartments in H3k27ac. b. The gene density in A/B compartment. c. The Gene Ontology (GO) analysis of TF in cluster1. d. The Gene Ontology (GO) analysis of TF in cluster2. e. The Gene Ontology (GO) analysis of TF target genes in cluster1. f. The Gene Ontology (GO) analysis of TF target genes in cluster2.

**Supplementary Figure 2.**
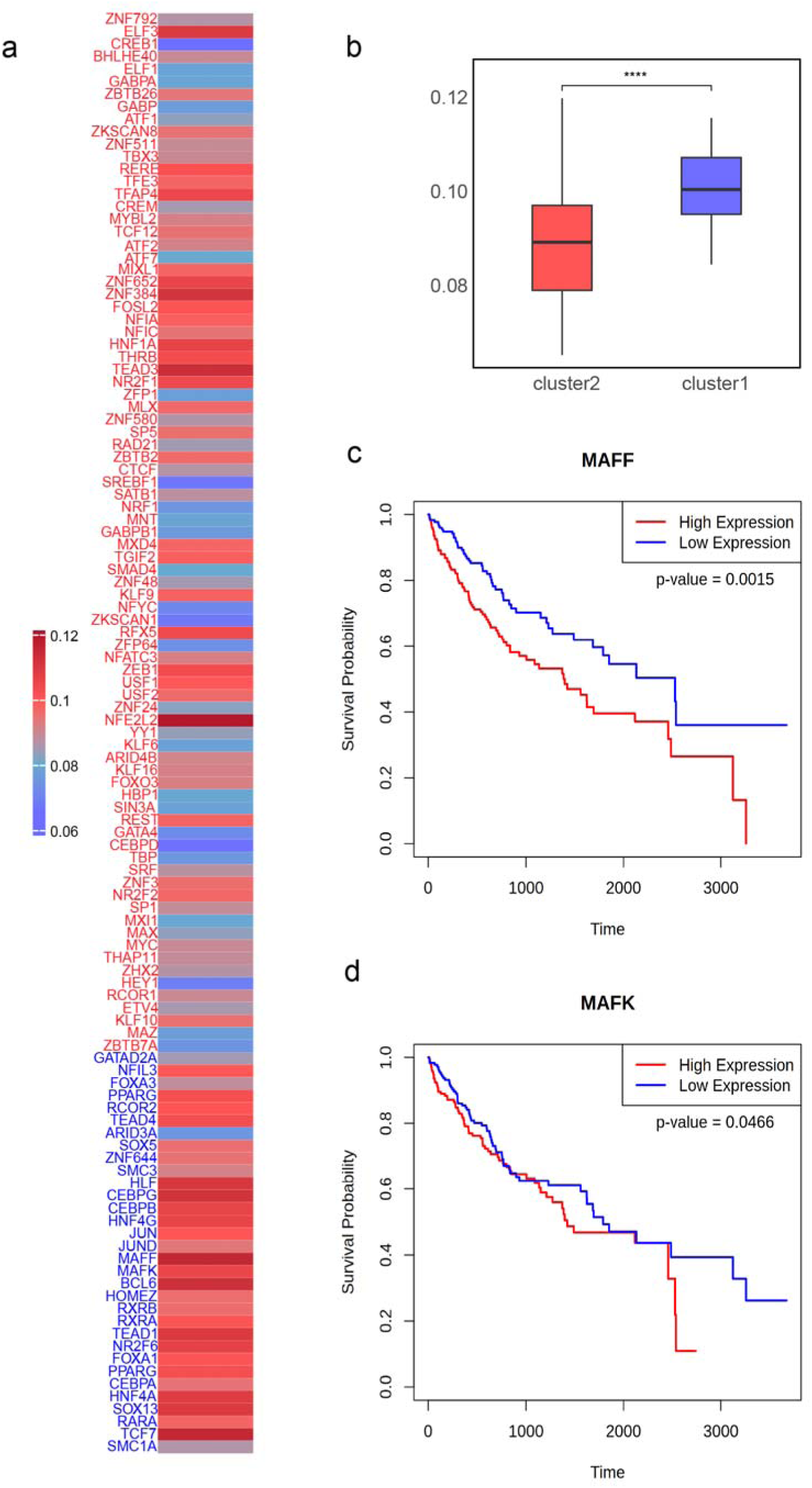
**a.** Overlap proportions between TF binding sites and eRNAs correlated with clinical outcomes, with darker red color indicating higher proportion of overlap. **b.** Comparison of overlap proportions between TF binding sites and eRNAs correlated with clinical outcomes across two distinct TF clusters. **c.** Survival analysis for MAFF. **d.** Survival analysis for MAFK.

**Supplementary Figure 3.**
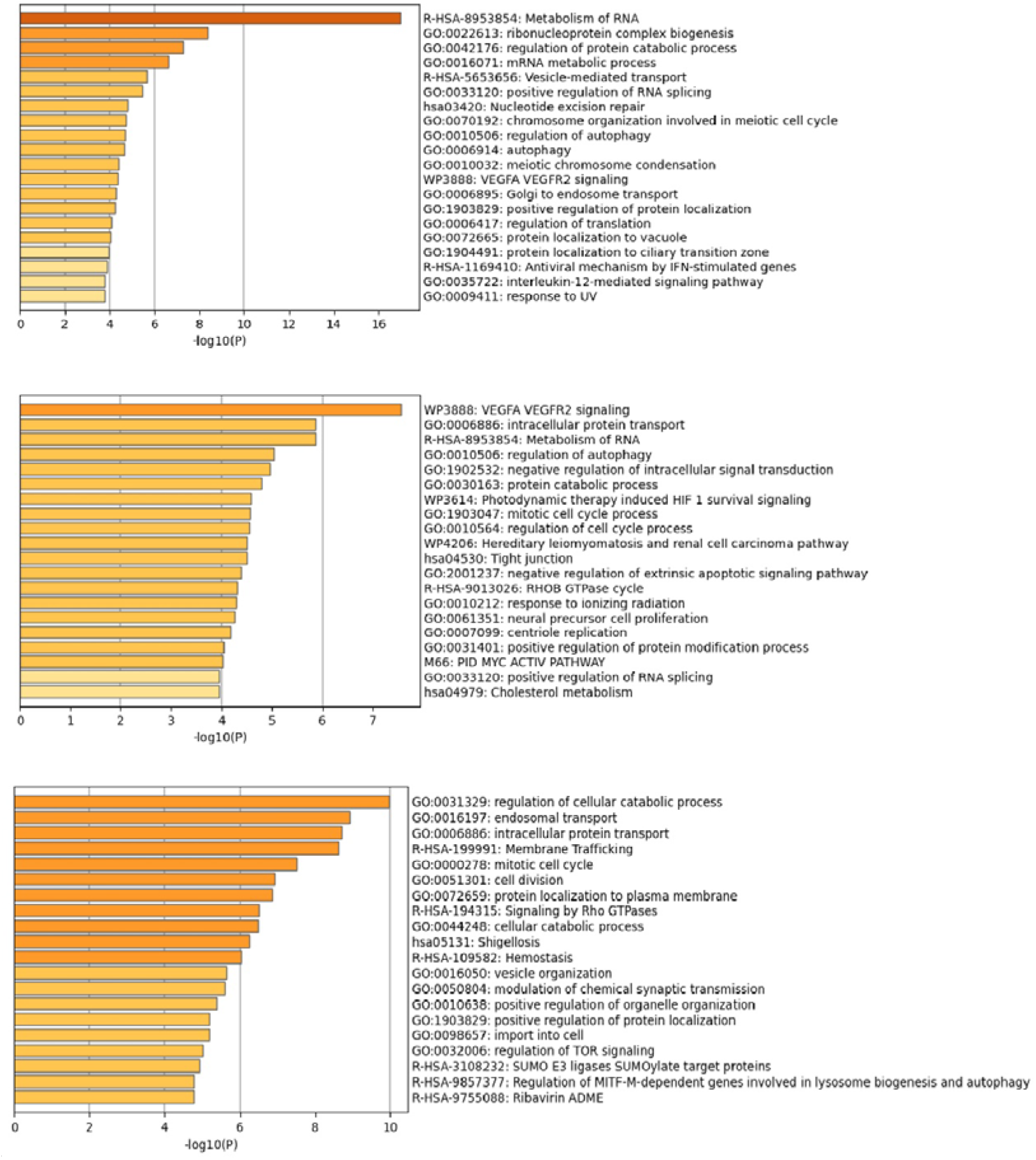
Gene Ontology (GO) analysis on the mRNAs interacting with top three eRNAs with the highest highest degrees;

**Supplementary Figure 4.**
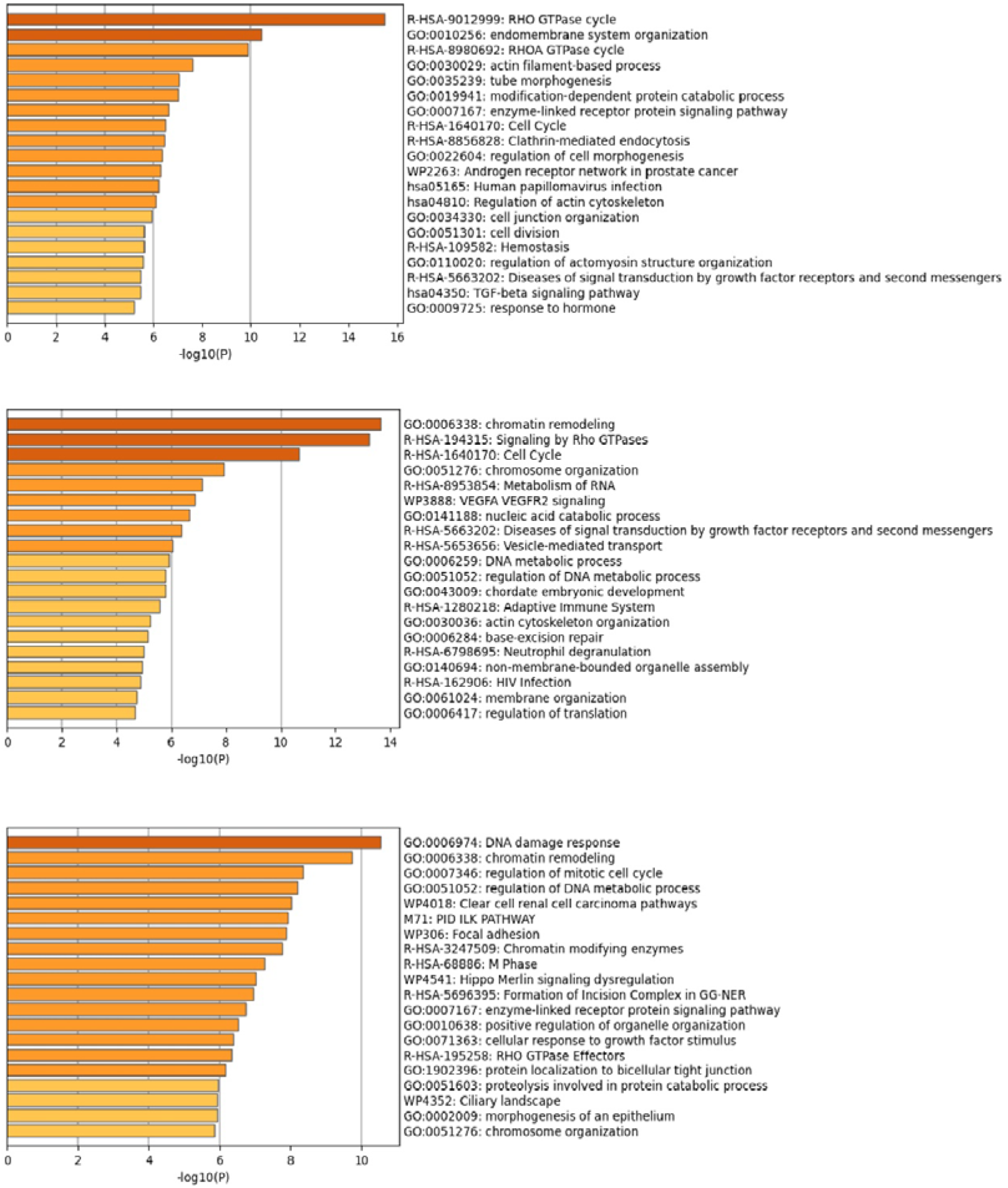
Gene Ontology (GO) analysis on the mRNAs interacting with top three uaRNAs with the highest highest degrees.

