## Supplementary_Methods for "Distinct RNA-RNA spatial interaction sub-networks of transcription factors in hepatic cells"

**Promoter-enhancer interaction(PEI) identification**

The ChIP–seq data of H3K4me3 and H3K27ac for hepG2 cell line were downloaded from the ENCODE^[^^1^^]^. The data were processed using Bowtie2(v2.4.4)^[^^2^^]^ with the following parameters: bowtie2 -p 16 -x GRCh38 -q H3k27ac.fastq -S H3k27ac.sam.The resulting SAM files were then sorted with Samtools(v1.6)^[^^3^^]^: samtools sort H3k27ac.sam -o H3k27ac.bam. Peak calling was performed using MACS2^[^^4^^]^ software with the following parameters: macs2 callpeak -f BAMPE -t H3k27ac.bam -c ControlS.bam -g hs -n H3k27ac --outdir macs2_result --nomodel -B -q 0.05 --extsize 146 --SPMR. Super-enhancers were defined based on the coverage of H3K27ac ChIP-seq signals using the software Rank Ordering of Super-Enhancers (ROSE)^[^^5^^]^.

We utilized both self-generated data (quality control in Supplementary Table1) and publicly available RIC-seq datasets of hepG2 cell line and carried out analyses to identify inter-molecular RNA-RNA interactions as previously described^[^^6^^]^. In brief, RIC-seq libraries were generated following the protocol described by Yuanchao Xue et al^[^^6^^]^. And adapter sequences were removed using the Trimmomatic program (v0.36)^[^^7^^]^, with the following parameters: java -jar trimmomatic-0.36.jar PE -phred33 -threads 10 fastq1.gz fastq2.gz read1.clean.pair.fq read1.clean.unpair.fq read2.clean.pair.fq read2.clean.unpair.fq ILLUMINACLIP:TruSeq3-PE-2.fa:2:30:7:8:true LEADING:25 TRAILING:20 SLIDINGWINDOW:4:15 MINLEN:30. Secondly, PCR duplicates were removed using custom scripts, and low-complexity fragments were trimmed from the ends of reads using Cutadapt (v1.15)^[^^8^^]^, with the following parameters: cutadapt -j 10 -b A{100} -b C{100} -b G{100} -b T{100} -n 3 --minimum-length=30 -e 0.1 -o -read.fq. Thirdly, reads were mapped to rRNA sequences using STAR software (v2.7.10a)^[^^9^^]^, with the following parameters: STAR --runMode alignReads --genomeDir index --readFilesIn read.fq --outFileNamePrefix prefix --outReadsUnmapped Fastx --outFilterMultimapNmax 100 --outSAMattributes All --outSAMtype BAM Unsorted --alignIntronMin 1 --scoreGapNoncan -4 --scoreGapATAC -4 --chimSegmentMin 15 --chimJunctionOverhangMin 15 --limitOutSJcollapsed 10000000 --limitIObufferSize 1500000000 1500000000 --runThreadN 20 --alignSJoverhangMin 15 --alignSJDBoverhangMin 10 --alignSJstitchMismatchNmax 5 -1 5 5. Fourthly, we constructed an index of the human reference genome and enhancer-promoter sequences. The remaining reads were aligned to the human reference genome using the STAR software, obtaining normally mapped reads (Aligned_out.sam) and chiastically mapped reads (Chimeric_out.sam). Intermolecular sequences were subsequently identified and extracted. Ultimately, high-confidence intermolecular RNA-RNA interactions were obtained.

Raw paired-end FASTQ

▼

Adapter trimming and quality filtering

└─ Trimmomatic

▼

PCR duplicate removal

└─ remove_duplicated_reads.pl

▼

Low-complexity sequence removal

└─ Cutadapt

▼

rRNA sequence mapping

└─ STAR

│

├───────────────┐

│	│

Mapped rRNA reads	Unmapped reads

│

▼

Reference genome mapping

└─ STAR

│

├───────────────────────┐

│	│

Conventional alignments	Chimeric alignments

│	│

▼	▼

Alignment filtering	Process Chimeric SAM

├─ MAPQ ≥ 30	└─ process_Chimeric_sam.pl

└─ Remove secondary alignments

│	│

└───────────┬───────────┘

▼

Identification of RNA-RNA interaction pairs

├─ Paired-end mapped reads

├─ Gapped/chimeric reads

└─ Merge all interaction

▼

All interaction pairs

▼

Separate intramolecular and intermolecular interactions

└ separate_intra_inter_pets.pl

▼

Intermolecular RNA-RNA interactions

**Speckle/ Nucleoli identification**

The interaction matrix for each pair of chromosomes (i, j) was generated the from Hi-C file using the Juicer^[^^10^^]^ command. The specific command used was: java -Xmx20g -jar juicer_tools.1.8.9_jCUDa.0.88.jar dump -d observed NONE hicfile.hic i j BP 500000 matrix_i_j.txt.

**ATAC-seq peak calling**

The downloaded ATAC-seq data were aligned to the genome using Bowtie2 with the following parameters: bowtie2 -p 8 -t -q -N 1 -L 25 -X 2000 --no-mixed --no-discordant -x index -1 myname1 -2 myname2 -S name.sam. Next, Samtools was used to remove alignments mapped to the mitochondrial genome with the following command: samtools view -hS {name}.sam | grep -v MT > name.rmmt.sam. The filtered alignments was then sorted using: samtools sort -@4 -m 1000000000 name.rmmt.rmq30.bam name.bam.sort. To remove duplicates, the following command was used: samtools rmdup name.bam.sort.bam name.rmdup.bam. Finally, MACS2 was employed to call peaks with the following command: macs2 callpeak –treatment=name.rmdup.bam.sort.bam –format=BAMPE --gsize=hs --outdir=output_dir --name=name.

**Loop identification**

To generate the loops, the Hi-C files were processed using the following command: java -Xms512m -Xmx20048m -Djava.library.path=lib_dir -jar Juicer_tools_path hiccups -m 500 -r 5000,10000 HiC_input -c chromosome output_file. The peak regions of transcription factors (TFs) were then overlapped with either end of the loops to assess their degree of enrichment.

**Clustering propensity (CP) score estimation**

We calculated CP score as previously described^[^^11^^]^. In brief, distribution of log10 transformed distances between each TF binding site and its nearest site was estimated, and two-sided K-S test was applied to compare the distribution of ChIP-seq binding profile (T) and that of the control (C), and obtained the CP score. 100 random sets of genomic intervals are generated as controls (C) to obtain 100 CP scores for each TF. The average of the 100 CPs was used as the final CP score. Bedtools was used to find the closest distance with parametres: bedtools closest - -a FILE1 -b FILE2 d-io -t first. The CP score is defined as follows:

A is defined as：Log_10_Distance (T) < Log_10_Distance (C).

B is defined as：Log_10_Distance (T) > Log_10_Distance (C).

CP is determined as

CP = A, A≥B

-B, A<B
